# Mitochondrial unfolded protein response transcription factor ATFS-1 increases resistance to exogenous stressors through upregulation of multiple stress response pathways

**DOI:** 10.1101/2021.04.26.441408

**Authors:** Sonja K. Soo, Annika Traa, Meeta Mistry, Jeremy M. Van Raamsdonk

## Abstract

The mitochondrial unfolded protein response (mitoUPR) is an evolutionarily conserved pathway that responds to various insults to the mitochondria through transcriptional changes that restore mitochondrial homeostasis in order to facilitate cell survival. Gene expression changes resulting from the activation of the mitoUPR are mediated by the transcription factor ATFS-1/ATF-5. To further define the mechanisms through which the mitoUPR protects the cell during mitochondrial dysfunction, we characterized the role of ATFS-1 in responding to organismal stress. We found that activation of ATFS-1 is sufficient to cause upregulation of genes involved in multiple stress response pathways, including the DAF-16-mediated stress response pathway, the SKN-1-mediated oxidative stress response pathway, the HIF-mediated hypoxia response pathway, the p38-mediated innate immune response pathway, and antioxidant genes. Moreover, ATFS-1 is required for the upregulation of stress response genes after exposure to exogenous stressors, especially oxidative stress and bacterial pathogens. Constitutive activation of ATFS-1 increases resistance to multiple acute exogenous stressors, while disruption of *atfs-1* decreases stress resistance. Although ATFS-1-dependent genes are upregulated in multiple long-lived mutants, constitutive activation of ATFS-1 in wild-type animals results in decreased lifespan. Overall, our work demonstrates that ATFS-1 serves a vital role in organismal survival of acute stresses through its ability to activate multiple stress response pathways, but that chronic ATFS-1 activation is detrimental for longevity.

## Introduction

The mitochondrial unfolded protein response (mitoUPR) is a stress response pathway that acts to reestablish mitochondrial homeostasis through inducing transcriptional changes of genes involved in metabolism and restoration of mitochondrial protein folding [1]. Various perturbations to the mitochondria can activate mitoUPR, including excess reactive oxygen species (ROS) and defects in mitochondrial import machinery [2]. The mitoUPR is mediated by the transcription factor ATFS-1 (activating transcription factor associated with stress-1) in *C. elegans* [3], or ATF5 in mammals [4].

ATFS-1/ATF5 regulates mitoUPR through its dual targeting domains, a mitochondrial targeting sequence (MTS) and a nuclear localization signal (NLS). Under normal unstressed conditions, the MTS causes ATFS-1 to enter the mitochondria through the HAF-1 import channel. Inside the mitochondria, ATFS-1 is degraded by the Lon protease CLPP-1/CLP1 [3]. However, mitochondrial stress disrupts ATFS-1 import into the mitochondria, resulting in cytoplasmic accumulation of ATFS-1. The NLS of the cytoplasmic ATFS-1 then targets it to the nucleus, where ATFS-1 acts with the transcription factor DVE-1 and transcriptional regulator UBL-5 to upregulate expression of chaperones, proteases, and other proteins [5].

In order to study the role of the mitoUPR in longevity, we previously disrupted *atfs-1* in long-lived *nuo-6* mutants, which contain a point mutation that affects Complex I of the electron transport chain [6]. *nuo-6* mutants have a mild impairment of mitochondrial function that leads to increased lifespan and enhanced resistance to multiple stressors. We found that loss of *atfs-1* not only decreased the lifespan of *nuo-6* worms, but also abolished the increased stress resistance of these worms, thereby suggesting that ATFS-1 contributes to both longevity and stress resistance in these worms [7].

While a role for the mitoUPR in longevity has been reported [8–11], and debated [12, 13], little is known about the role of ATFS-1 in response to exogenous stressors. Pellegrino *et al*. found that activation of ATFS-1 can increase organismal resistance to the pathogenic bacteria *P. aeruginosa* [14], while Pena *et al*. showed that ATFS-1 activation can protect against anoxia-reperfusion-induced death [15].

In this study, we use *C. elegans* to define the relationship between ATFS-1 and organismal stress resistance, and explore the underlying mechanisms. We find that activation of ATFS-1 is sufficient to upregulate genes from multiple stress response pathways and is important for transcriptional changes induced by oxidative stress and bacterial pathogen exposure. Constitutive activation of ATFS-1 is also sufficient to increase resistance to multiple stressors. While ATFS-1-dependent genes are upregulated in several long-lived mutants representative of multiple pathways of lifespan extension, chronic activation of ATFS-1 does not extend longevity. Overall, our results demonstrate a crucial role for ATFS-1 in organismal stress response through activation of multiple stress response pathways.

## Results

### ATFS-1 activates genes from multiple stress response pathways

Mild impairment of mitochondrial function through a mutation in *nuo-6* results in the activation of the mitoUPR. We previously performed a bioinformatic analysis of genes that are upregulated in *nuo-6* mutants in an ATFS-1-dependent manner, and discovered an enrichment for genes associated with the GO term “response to stress” [7]. Based on this observation, we hypothesized that ATFS-1 may be able to activate other stress response pathways. To test this hypothesis, we quantified the expression of established target genes from eight different stress response pathways under conditions where ATFS-1 is either activated, or where ATFS-1 is disrupted.

To activate ATFS-1, we used the *nuo-6* mutation. We also examined gene expression in two different gain-of-function (GOF) mutants with constitutively active ATFS-1: *atfs-1(et15)* and *atfs-1(et17)*. Both of these constitutively active ATFS-1 mutants have mutations in the MTS causing increased nuclear localization of ATFS-1 [16]. We used a loss-of-function (LOF) *atfs-1* deletion mutation (*gk3094*) to disrupt ATFS-1 function in wild-type and *nuo-6* mutants.

Expression of specific stress response target genes were examined: *hsp-6* in the mitochondrial unfolded protein response (mitoUPR) pathway; *hsp-4* in the endoplasmic reticulum unfolded protein response (ER-UPR) pathway; *hsp-16.2* in the cytoplasmic unfolded protein response pathway (cytoUPR); *sod-3* in the DAF-16-mediated stress response pathway; *gst-4* in the SKN-1-mediated stress response pathway; *nhr-57* in the HIF-1-mediated hypoxia response pathway; *Y9C9A.8* in the p38-mediated innate immunity pathway; and *trx-2*, an antioxidant gene (**Table S1**).

We found that compared to wild-type worms, *atfs-1(gk3094)* deletion mutants did not have decreased expression levels for the target genes of any of the stress response pathways (**Fig. 1**). This indicates that ATFS-1 is not required for the basal expression levels of these stress response genes.

**Figure 1.**
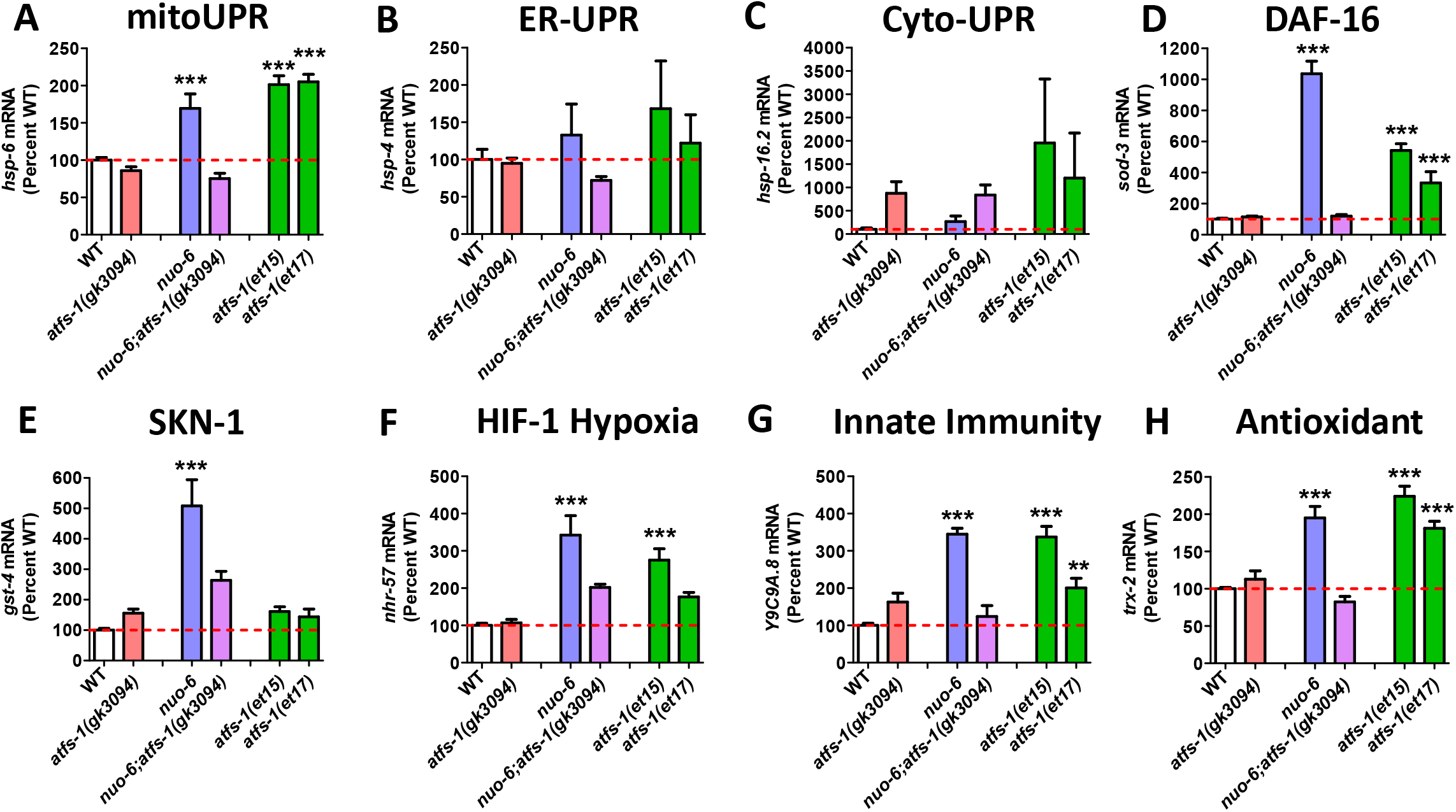
Activation of ATFS-1 upregulates genes from multiple stress response pathways. To determine the role of ATFS-1 in the activation of genes from different stress response pathways, we activated ATFS-1 by mildly impairing mitochondrial function through a mutation in *nuo-6* (blue bars) and then examined the effect of disrupting *atfs-1* using an *atfs-1* deletion mutant *atfs-1(gk3094)* (purple bars). We also examined the expression of these genes in two constitutively active *atfs-1* mutants, *atfs-1(et15)* and *atfs-1(et17)* (green bars). Target genes from the mitochondrial unfolded protein response (mitoUPR) (**A**, *hsp-6*), DAF-16-mediated stress response (**D**, *sod-3*), SKN-1-mediated oxidative stress response (**E**, *gst-4*), HIF-1-mediated hypoxia response (**F**, *nhr-57*), p38-mediated innate immune pathway (**G**, *Y9C9A.8*), and antioxidant defense (**H**, *trx-2*) are all significantly upregulated in *nuo-6* mutants in an ATFS-1-dependent manner. Target genes from the mitoUPR, DAF-16-mediated stress response, HIF-1-mediated hypoxia response, p38-mediated innate immune pathway and antioxidant defense are also upregulated in constitutive activation of ATFS-1 (**A, D, F, G, H**). In contrast, activation of ATFS-1 by *nuo-6* mutation or *atfs-1* gain-of-function mutations did not significantly affect target gene expression for the endoplasmic reticulum unfolded protein response (**B**, ER-UPR, *hsp-4*) or the cytoplasmic unfolded protein response (**C**, Cyto-UPR, *hsp-16.2*). *atfs-1(gk3094)* is a loss-of-function deletion mutant. *atfs-1(et15)* and *atfs-1(et17)* are constitutively active gain-of-function mutants. Error bars indicate SEM. **p<0.01, ***p<0.001. A full list of genes that are upregulated by ATFS-1 activation can be found in **Table S2**.

Activation of the mitoUPR through mutation of *nuo-6* resulted in significant upregulation of target genes from the mitoUPR (*hsp-6*; **Fig. 1A**), the DAF-16-mediated stress response (*sod-3*; **Fig. 1D**), the SKN-1-mediated oxidative stress response (*gst-4*; **Fig. 1E**), the HIF-1-mediated hypoxia response (*nhr-57*; **Fig. 1F**), the p38-mediated innate immunity pathway (*Y9C9A.8*; **Fig. 1G**), and antioxidant defense (*trx-2*; **Fig. 1H**). Importantly, for all of these genes, inhibiting the mitoUPR through deletion of *atfs-1* prevented the upregulation of the stress response in *nuo-6;atfs-1(gk3094)* worms (**Fig. 1A, D-H**), indicating that ATFS-1 is required for the activation of these stress pathway genes during mitochondrial stress.

Constitutive activation of ATFS-1 in *atfs-1(et 15)* and *atfs-1(et17)* mutants resulted in upregulation of most of the same genes that are upregulated in *nuo-6* mutants, except for *gst-4* in the SKN-1 pathway (**Fig. 1A, D-H**). This indicates that ATFS-1 activation is sufficient to induce upregulation of specific stress response genes independently of mitochondrial stress. Activating the mitoUPR through the *nuo-6* mutation, or through the constitutively-active ATFS-1 mutants did not significantly increase the expression of target genes from the ER-UPR (*hsp-4*; **Fig. 1B**) or the cyto-UPR (*hsp-16.2*; **Fig. 1C**).

To gain a more comprehensive view of the extent to which mitoUPR activation causes upregulation of genes in other stress response pathways, we compared genes upregulated in the constitutively active *atfs-1* mutant, *atfs-1(et15)*, to genes upregulated by activation of different stress response pathways. As a proof-of-principle, we first examined the overlap between upregulated genes in *atfs-1(et15)* mutants and genes upregulated by activation of the mitoUPR with *spg-7* RNAi in an ATFS-1-dependent manner [3].

We identified genes upregulated by the activation of other stress response pathways from published gene expression studies, and the genes and relevant pathways are listed in **Table S3**. Target genes from the ER-UPR pathway were defined as genes upregulated by tunicamycin exposure and dependent on either *ire-1*, *xbp-1*, *pek-1* or *atf-6* [17]. Cyto-UPR pathway genes are genes upregulated by overexpression of heat shock factor 1 (HSF-1) and genes bound by HSF-1 after a thirty-minute heat shock at 34°C [18, 19]. DAF-16 pathway genes were identified by Tepper et al. by performing a meta-analysis of 46 previous gene expression studies, comparing conditions in which DAF-16 is activated (e.g. *daf-2* mutants) and conditions in which the activation is inhibited by disruption of *daf-16* (e.g. *daf-2;daf-16* mutants) [20]. SKN-1 pathway genes were identified as genes that exhibit decreased expression after *skn-1* RNAi in wild-type worms, genes that are upregulated in *glp-1* mutants in a SKN-1-dependent manner, genes that are upregulated by germline stem cell removal in a SKN-1-dependent manner [21], and genes upregulated in *daf-2* mutants in a SKN-1-dependent manner [22]. HIF-1-mediated hypoxia genes are genes induced by hypoxia in a HIF-1-dependent manner [23]. Innate immunity genes are defined as genes upregulated by exposure to *Pseudomonas aeruginosa* strain PA14 in a PMK-1- and ATF-7-dependent manner [24], where PMK-1 and ATF-7 are part of the p38-mediated innate immune signaling pathway. Finally, antioxidant genes is a comprehensive list of genes involved in antioxidant defense such as superoxide dismutases (*sod*), catalases (*ctl*), peroxiredoxins (*prdx*), or thioredoxins (*trx*).

In comparing genes upregulated in the constitutively active *atfs-1* mutant *et15* to these previously published gene lists, we found that 51% of genes upregulated by *spg-7* RNAi in an ATFS-1-dependent manner are also upregulated by constitutive activation of ATFS-1 (**Fig. 2A**). Similarly, we found a highly significant overlap of upregulated genes between *atfs-1(et15)* mutants and each of the examined stress response pathways. We found that *atfs-1(et15)* had a 25% overlap with genes of ER-UPR pathway (**Fig. 2B**); 22% overlap with genes of the Cyto-UPR pathway (**Fig. 2C**); 26% overlap with genes of the DAF-16-mediated stress pathway (**Fig. 2D**); 30% overlap with genes of the SKN-1-mediated stress pathway (**Fig. 2E**); 23% overlap with genes of the HIF-1-mediated hypoxia pathway (**Fig. 2F**); 22% overlap with genes of the p38-mediated innate immunity pathway (**Fig. 2G**); and 33% overlap with antioxidant genes (**Fig. 2H**). Combined, this indicates that activation of ATFS-1 is sufficient to upregulate genes in multiple stress response pathways.

**Figure 2.**
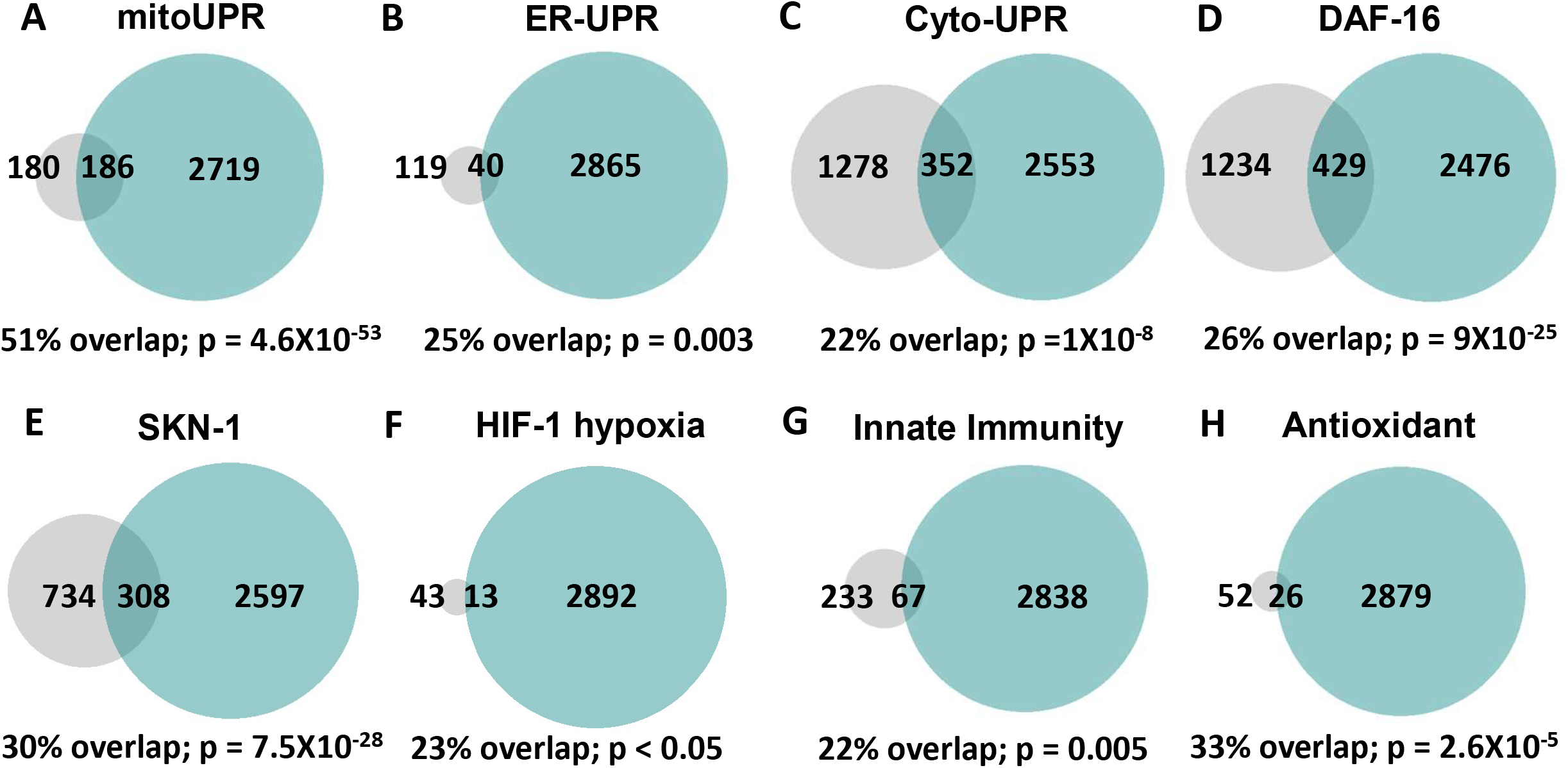
Constitutive activation of ATFS-1 results in upregulation of genes from multiple stress response pathways. Genes that are upregulated by activation of ATFS-1 were compared to previous published lists of genes involved in different stress response pathways, including the mitochondrial unfolded protein response (**A**, mitoUPR), the endoplasmic reticulum unfolded protein response (**B**, ER-UPR), the cytoplasmic unfolded protein response (**C**, Cyto-UPR), the DAF-16-mediated stress response (**D**), the SKN-1-mediated oxidative stress response (**E**), the HIF-1-mediated hypoxia response (**F**), the p38-mediated innate immune response (**G**), and antioxidant genes (**H**). In every case, there was a significant degree of overlap ranging from 22%-51%. Grey circles indicate genes that are upregulated by activation of the stress response pathway indicated. Turquoise circles indicate genes that are upregulated in the *atfs-1(et15)* constitutively active gain-of-function mutant. The numbers inside the circles show how many genes are upregulated. The percentage overlap is the number of overlapping genes as a percentage of the number of genes upregulated by the stress response pathway. mitoUPR = mitochondrial unfolded protein response. ER-UPR = endoplasmic reticulum unfolded protein response. Cyto-UPR = cytoplasmic unfolded protein response. DAF-16 = DAF-16-mediated stress response pathway. SKN-1 = SKN-1-mediated oxidative stress response pathway. HIF-1 = HIF-1-mediated stress response pathway. Innate immunity = p38-mediated innate immunity pathway. Antioxidant = antioxidant genes. Stress pathway gene lists and sources can be found in **Table S3**.

### ATFS-1 is required for transcriptional responses to exogenous stressors

Having shown that constitutive activation of ATFS-1 can induce upregulation of genes involved in various stress response pathways, we next sought to determine the role of ATFS-1 in the genetic response to different stressors. To do this, we exposed wild-type animals and *atfs-1* loss-of-function mutants (*atfs-1(gk3094)*) to six different types of stress and quantified the resulting upregulation of stress response genes using quantitative RT-PCR (qPCR). We found that exposure to either oxidative stress (4 mM paraquat, 48 hours) or the bacterial pathogen *Pseudomonas aeruginosa* strain PA14 induced a significant upregulation of stress response genes in wild-type worms, but that this upregulation was suppressed by disruption of *atfs-1* (**Fig. 3A,B**). In contrast, exposure to heat stress (35°C, 2 hours; **Fig. 3C**), osmotic stress (300 mM NaCl, 24 hours; **Fig. 3D**), anoxic stress (24 hours; **Fig. 3E**), or ER stress (tunicamycin for 24 hours; **Fig. 3F**) caused upregulation of stress response genes in both wild-type and *atfs-1(gk3094)* worms to a similar extent. Combined, these results indicate that ATFS-1 is required for upregulating stress response genes in response to exposure to oxidative stress or bacterial pathogens.

**Figure 3.**
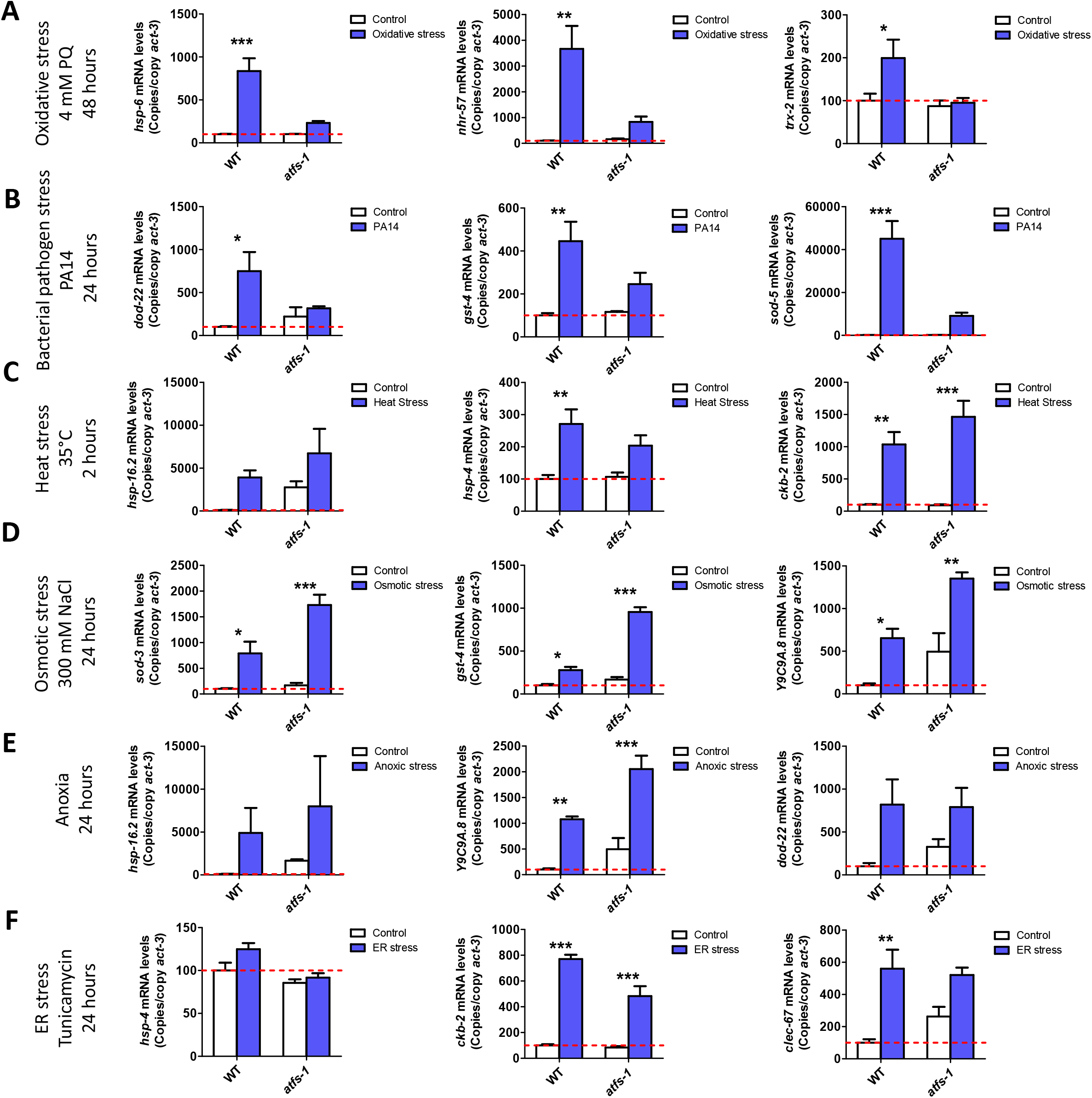
ATFS-1 is required for upregulation of stress response genes after exposure to oxidative stress or bacterial pathogen stress. To determine the role of ATFS-1 in responding to different types of stress, we compared the upregulation of stress response genes in wild-type and *atfs-1(gk3094)* loss-of-function deletion mutants after exposure to different stressors. **A.** Exposure to oxidative stress (4 mM paraquat, 48 hours) caused a significant upregulation of *hsp-6*, *nhr-57* and *trx-2* in wild-type worms that was prevented by the disruption of *atfs-1*. **B.** Exposure to bacterial pathogen stress (PA14, 24 hours) resulted in an upregulation of *dod-22*, *gst-4* and *sod-5* in wild-type worms that was prevented by the *atfs-1* deletion. **C.** Exposure to heat stress (35°C, 2 hours) increased the expression of *hsp-16.2*, *hsp-4* and *ckb-2* in both wildtype and *atfs-1* worms. **D.** Exposure to osmotic stress (300 mM, 24 hours) caused an upregulation of *sod-3*, *gst-4* and *Y9C9A.8* in wild-type worms and to a greater magnitude in *atfs-1* mutants. **E.** Anoxia (24 hours) resulted in the upregulation of *hsp-16.2*, *Y9C9A.8* and *dod-22* in both wild-type and *atsf-1* worms. **F.** Exposing worms to endoplasmic reticulum stress (tunicamycin, 24 hours) increased the expression of *ckb-2* and *clec-67* in both wild-type and *atfs-1* worms. Error bars indicate SEM. *p<0.05, **p<0.01, ***p<0.001.

### Modulation of ATFS-1 levels affects resistance to multiple stressors

Due to the crucial role of ATFS-1 in upregulating genes in multiple stress response pathways, we next sought to determine the extent to which activating ATFS-1 protects against exogenous stressors. To do this, we quantified resistance to stress in two constitutively active *atfs-1* gain-of-function mutants (*atfs-1(et15)*, *atfs-1(et17)*) compared to wild-type worms. For comparison, we also included an *atfs-1* loss-of-function deletion mutant (*atfs-1(gk3094)*), which we have previously shown to have decreased resistance to oxidative, heat, osmotic and anoxic stress [7].

We measured resistance to acute oxidative stress by exposing worms to 300 μM juglone. We found that both the gain-of-function mutants, *atfs-1(et15)* and *atfs-1(et17)*, have increased resistance to acute oxidative stress compared to wild-type worms, while *atfs-1(gk3094)* deletion mutants were less resistant compared to wild-type worms (**Fig. 4A**). To test resistance to chronic oxidative stress, worms were transferred to plates containing 4 mM paraquat beginning at day 1 of adulthood. Similar to the acute assay, *atfs-1(et17)* mutants were more resistant to chronic oxidative stress, while *atfs-1(gk3094)* mutants were less resistant to chronic oxidative stress compared to wild-type worms (**Fig. 4B**). Oddly, *atfs-1(et15)* gain-of-function mutants exhibited decreased resistance to chronic oxidative stress.

**Figure 4.**
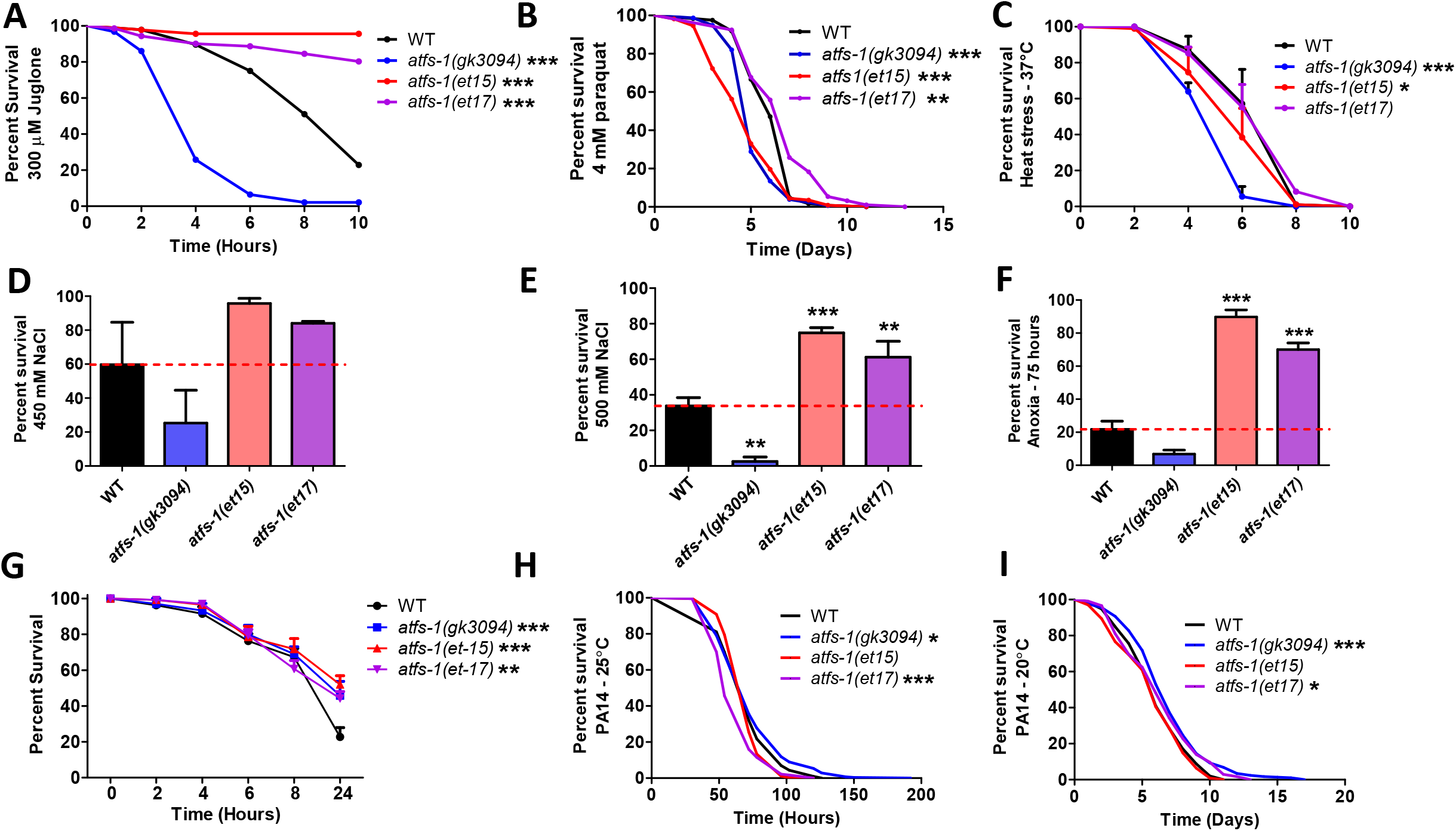
Constitutive activation of ATFS-1 increases resistance to multiple external stressors. To determine the role of ATFS-1 in resistance to stress, the stress resistance of an *atfs-1* loss-of-function mutants (*atfs-1(gk3094)*) and two constitutively active *atfs-1* gain-of-function mutants (*atfs-1(et15)*, *atfs-1(et17)*) was compared to wild-type worms. **A.** Activation of ATFS-1 enhanced resistance to acute oxidative stress (300 μM juglone), while deletion of *atfs-1* markedly decreased resistance to acute oxidative stress. **B.** Disruption of *atfs-1* decreased resistance to chronic oxidative stress (4 mM paraquat). *atfs-1(et17)* mutants showed increased resistance to chronic oxidative stress, while *atfs-1(et15)* mutants had decreased resistance. **C.** Resistance to heat stress (37°C) was not enhanced by activation of ATFS-1, while deletion of *atfs-1* decreased heat stress resistance. **D,E.** Activation of ATFS-1 increased resistance to osmotic stress (450 mM, 500 mM NaCl), while disruption of *atfs-1* decreased osmotic stress resistance. **F.** Constitutively active *atfs-1* mutants show increased resistance to anoxia (75 hours), while *atfs-1* deletion mutants exhibit a trend towards decreased anoxia resistance. **G.** Activation of ATFS-1 increased resistance to *P. aeruginosa* toxin in a fast kill assay. A slow kill assay in which worms die from internal accumulation of *P aeruginosa* was performed according to two established protocols. **H.** At 25°C, *atfs-1(et17)* mutants showed a small decrease in resistance to bacterial pathogens (PA14), while *atfs-1(gk3094)* mutants showed a small increase in resistance. **I.** At 20°C, both *atfs-1(et17)* and *atfs-1(gk3094)* mutants exhibited a small increase in resistance to bacterial pathogens. Error bars indicate SEM. *p<0.05, **p<0.01, ***p<0.001.

Resistance to heat stress was measured at 37°C. None of the mutants showed increased survival during heat stress, with *atfs-1(et15)* and *atfs-1(gk3094)* mutants both showing a significant decrease in survival compared to wild-type worms (**Fig. 4C**). Resistance to osmotic stress was quantified on plates containing 450 mM or 500 mM NaCl after 48 hours. At both concentrations, the constitutively active *atfs-1* mutants had increased survival compared to wild-type worms, while *atfs-1(gk3094)* deletion mutants had decreased survival, although the difference was only significant at 500 mM (**Fig. 4D, E**). Resistance to anoxic stress was measured by placing worms in an oxygen-free environment for 75 hours, followed by a 24-hour recovery. We observed increased survival in *atfs-1(et15)* and *atfs-1(et17)* mutants and decreased survival in *atfs-1(gk3094)* mutant compared to wild-type worms (**Fig. 4F**).

Lastly, to test resistance to bacterial pathogens, worms were exposed to *Pseudomonas aeruginosa* strain PA14 in either a fast kill assay in which worms die from a toxin produced by the bacteria, or a slow kill assay in which worms die due to the intestinal colonization of the pathogenic bacteria [25]. In the fast kill assay, we found that constitutive activation of ATFS-1 increases survival in *atfs-1(et15)* and *atfs-1(et17)* mutants (**Fig. 4G**). We also observed increased survival in *atfs-1(gk3094)* deletion mutants. For the slow kill assay, we used two established protocols: one in which the assay is initiated at the L4 larval stage and performed at 25°C [14, 25, 26] and one in which the assay is initiated at day three of adulthood and performed at 20°C [27]. Surprisingly, at 25°C, we found that *atfs-1(et17)* mutant had a small decrease in resistance to PA14, while *atfs-1(gk3094)* mutants exhibited a small increase in resistance to PA14 compared to wild-type worms (**Fig. 4H**). At 20°C, both *atfs-1(gk3094)* and *atfs-1(et17)* mutants had a small increase in resistance to PA14 compared to wild-type worms (**Fig. I**).

All together, these data indicate that activation of ATFS-1 is sufficient to protect against oxidative stress, osmotic stress, anoxia, and bacterial pathogens but not heat stress. They also show that ATFS-1 is required for wild-type worms to survive oxidative stress, heat stress, osmotic stress, and anoxia.

### Long-lived genetic mutants have upregulation of ATFS-1 target genes

We previously showed that ATFS-1 target genes are upregulated in three long-lived mitochondrial mutants: *clk-1*, *isp-1* and *nuo-6* [6, 7, 28, 29]. To determine if ATFS-1 target genes are specifically upregulated in long-lived mitochondrial mutants, or if they are also upregulated in other mutants with extended longevity, we compared gene expression in six additional long-lived mutants, which act through other longevity-promoting pathways, to genes that are upregulated by ATFS-1 activation. These long-lived mutants included *sod-2* mutants, which act through increasing mitochondrial reactive oxygen species (ROS) [30]; *daf-2* mutants, which have reduced insulin-IGF1 signaling [31]; *glp-1* mutants, which have germline ablation [32]; *ife-2* mutants, which have reduced translation [33]; *osm-5* with reduced chemosensation [34]; and *eat-2* with dietary restriction [35].

After identifying genes that are differentially expressed in each of these long-lived mutants, we compared the differentially expressed genes to genes that are upregulated by ATFS-1 activation. We defined ATFS-1-upregulated genes in two ways: (1) genes that are upregulated by *spg-7* RNAi in an ATFS-1-dependent manner [3]; and (2) genes that are upregulated in a constitutively active *atfs-1* mutant (*et15*; [7]).

We found that the majority of the long-lived mutants examined had a significant enrichment of ATFS-1 target genes. In comparing the number of overlapping genes between genes upregulated in each long-lived mutant and genes upregulated by *spg-7* RNAi in an ATFS-1-dependent manner, the degree of overlap was significantly greater than would be expected by chance for *clk-1* (6.7 fold enrichment), *isp-1* (6.0 fold enrichment), *sod-2* (5.5 fold enrichment), *nuo-6* (4.1 fold enrichment), *daf-2* (2.6 fold enrichment), *glp-1* (2.0 fold enrichment), and *ife-2* mutants (1.5 fold enrichment)(**Fig. 5**). We did not find a significant enrichment of ATFS-1 targets in *osm-5* and *eat-2* worms (**Fig. 5**).

**Figure 5.**
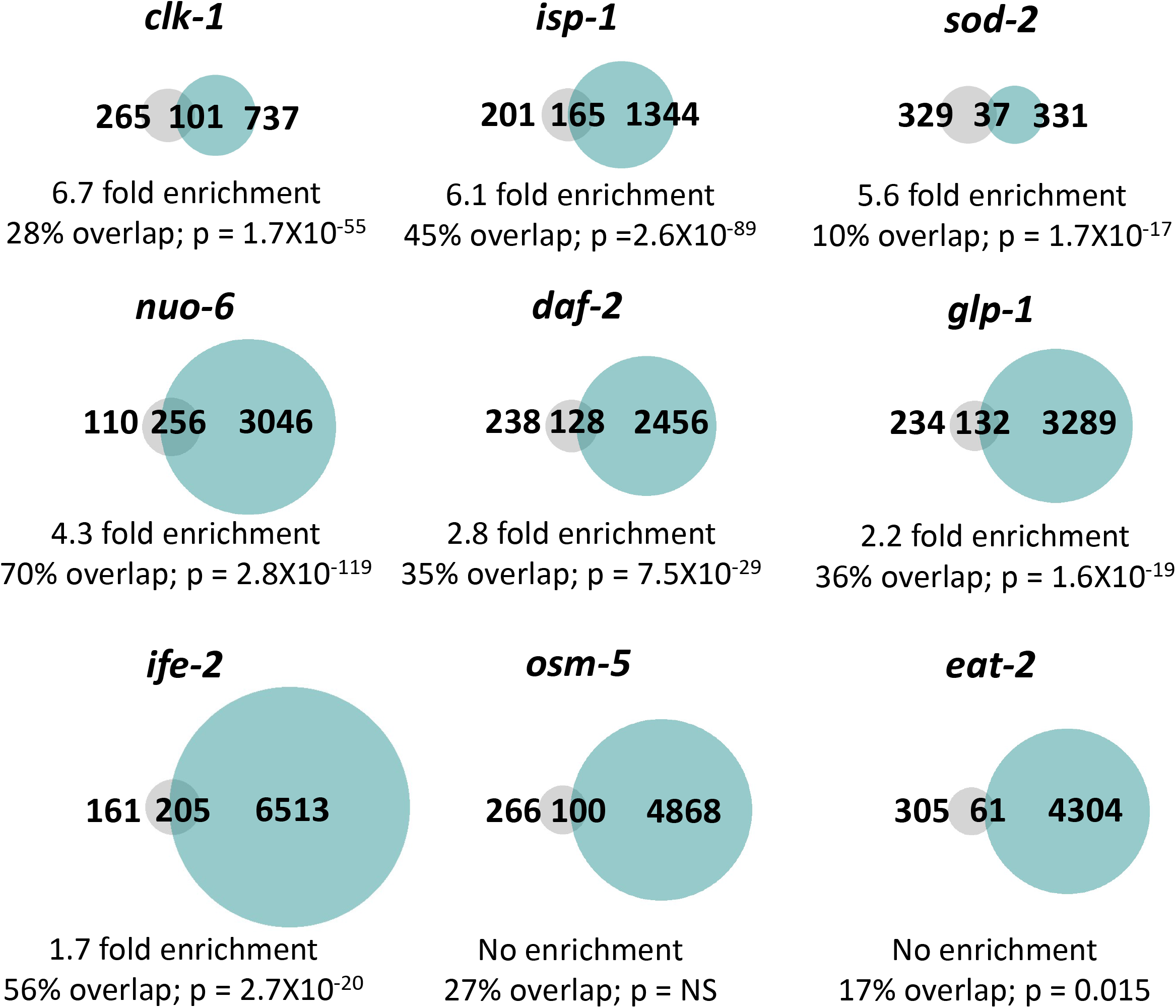
Multiple long-lived mutants from different pathways of lifespan extension show upregulation of ATFS-1-dependent genes. To determine the extent to which long-lived genetic mutants from different pathways of lifespan extension show differential expression of ATFS-1 target genes, we compared genes that are upregulated in nine different long-lived mutants to a published list of *spg-7* RNAi-upregulated, ATFS-1-dependent target genes (Nargund *et al*., 2012). We found that *clk-1*, *isp-1*, *nuo-6*, *sod-2*, *daf-2*, *glp-1* and *ife-2* worms all show a highly significant degree of overlap with genes upregulated by *spg-7* RNAi in an ATFS-1-dependent manner. The grey circles represent the 366 genes that are upregulated by *spg-7* RNAi in an ATFS-1 dependent manner. Turquoise circles are genes that are significantly upregulated in the long-lived mutant indicated as determined from our RNA sequencing data. The number of unique and overlapping genes are indicated. Overlap is calculated as the number of genes in common between the two gene sets divided by the total number of genes that are upregulated by *spg-7* RNAi in an ATFS-1 dependent manner. Enrichment is calculated as the number of overlapping genes observed divided by the number of overlapping genes predicted if genes were chosen randomly.

In comparing the number of overlapping genes between genes upregulated in each long-lived mutant and genes upregulated in the constitutively active *atfs-1(et15)* mutant, we found that the degree of overlap was significantly greater than would be expected by chance for *isp-1* (3.5 fold enrichment), *sod-2* (3.4 fold enrichment), *clk-1* (3.3 fold enrichment), *nuo-6* (2.5 fold enrichment), *daf-2* (2.4 fold enrichment), *glp-1* (1.8 fold enrichment), *ife-2* (1.8 fold enrichment), and *eat-2* mutants (1.5 fold enrichment) (**Fig. S1**). We did not observe a significant enrichment of ATFS-1 targets in *osm-5* mutants (**Fig. S1**).

Overall, these results indicate that ATFS-1 target genes are upregulated in multiple long-lived mutants, including mutants in which mitochondrial function is not directly disrupted.

### Constitutively active *atfs-1* mutants have decreased lifespan despite enhanced resistance to stress

Having shown that ATFS-1 target genes are activated in multiple long-lived mutants, we sought to determine if ATFS-1 activation is sufficient to increase lifespan, and whether the presence of ATFS-1 is required for normal longevity in wild-type worms. Despite having increased resistance to multiple stresses, both constitutively active *atfs-1* mutants (*et15* and *et17*) have decreased lifespan compared to wild-type worms (**Fig. 6A,B**), which is consistent with a previous study finding shortened lifespan in *atfs-1(et17)* and *atfs-1(et18)* worms [13]. Despite having decreased resistance to multiple stresses, *atfs-1* deletion mutants (*gk3094*), had a lifespan comparable to wild-type worms (**Fig. 6C**), as we have previously observed [7]. Combined, this indicates that ATFS-1 does not play a major role in lifespan determination in a wild-type background despite having an important role in stress resistance.

**Figure 6.**
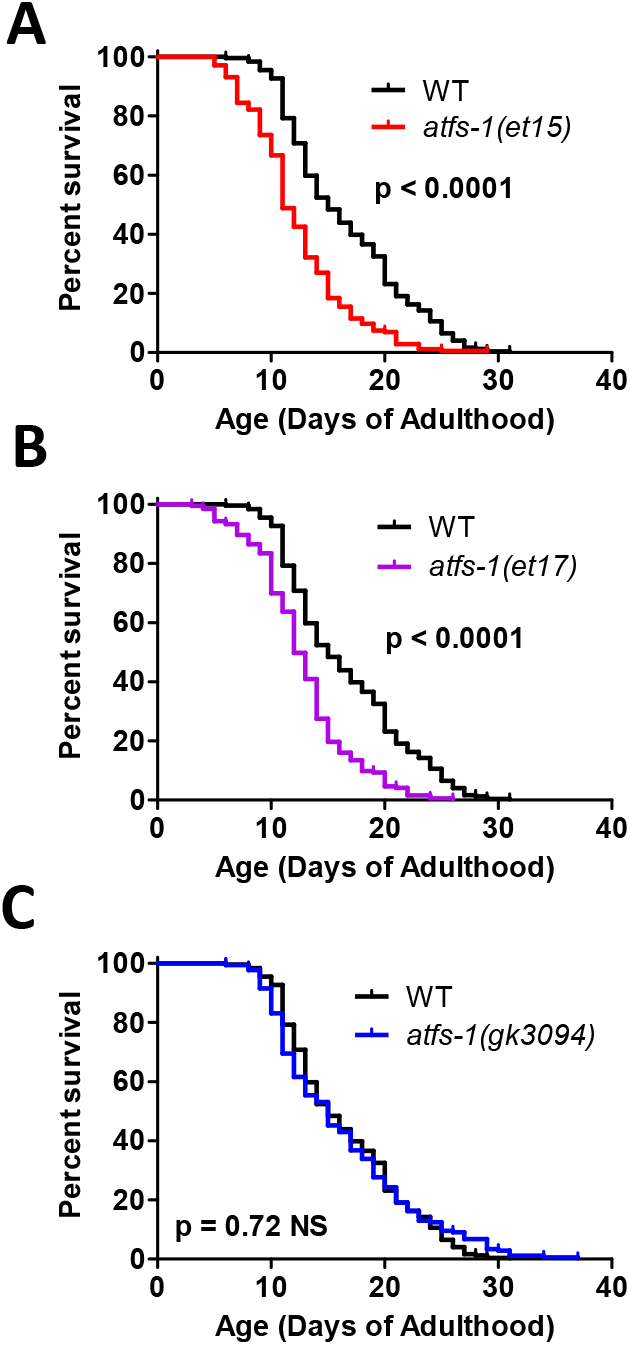
Activation of ATFS-1 does not increase lifespan. To determine the effect of ATFS-1 on aging, we quantified the lifespan of an *atfs-1* deletion mutant and two constitutively active *atfs-1* mutants. **A,B.** Both constitutively active *atfs-1* mutants, *et15* and *et17*, have a significantly decreased lifespan compared to wild-type worms. **C.** Deletion of *atfs-1* does not affect lifespan compared to wild-type worms. *atfs-1(gk3094)* is a loss of function mutant resulting from a deletion. *atfs-1(et15)* and *atfs-1(et17)* are constitutively active gain-of-function mutants.

## Discussion

Mitochondria are vital for organismal health as they perform multiple crucial functions within the cell including energy generation, metabolic reactions and intracellular signaling. Accordingly, it is important for cell and organismal survival maintain mitochondrial function during times of acute stress, and throughout normal aging. The mitoUPR is a conserved pathway that facilitates restoration of mitochondrial homeostasis after internal or external stresses. In this work, we demonstrate a crucial role for the mitoUPR transcription factor ATFS-1 in the genetic response to external stressors which ultimately promotes survival of the organism.

### ATFS-1 is not required for normal longevity

A number of studies have directly or indirectly examined the role of the mitoUPR and or ATFS-1 in longevity. In these studies, mitoUPR activation was typically measured using a mitoUPR reporter strain that expresses GFP under the promoter of *hsp-6*, which is a target gene of ATFS-1 and the mitoUPR.

A relationship between the mitoUPR and longevity was initially supported by the observation that disruption of the mitochondrial electron transport chain (ETC) by RNAi knockdown of the <u>c</u>ytochrome <u>c</u> <u>o</u>xidase-1 (*cco-1*) gene resulted in both increased lifespan [36] and activation of the mitoUPR [8, 37]. Since then, other lifespan-extending mutations have also been shown to activate the mitoUPR, including three long-lived mitochondrial mutants, *clk-1*, *isp-1* and *nuo-6* [7].

To explore this relationship in more comprehensive manner, Runkel *et al*. compiled a list of genes that activate the mitoUPR and looked at their effect on lifespan. Of the 99 genes reported to activated the mitoUPR, 58 resulted in increased lifespan, while only 7 resulted in decreased lifespan [38]. Bennet *et al*. performed an RNAi screen to identify RNAi clones that increase expression of a reporter of mitoUPR activity (*hsp-6p::GFP*) and measured the effect of a selection of these clones on lifespan [13]. Of the 19 mitoUPR-inducing RNAi clones that they tested, 10 RNAi clones increased lifespan, while 6 decreased lifespan [13]. Using a similar approach to screen for compounds that activate a mitoUPR reporter strain (*hsp-6p::GFP*), metolazone was identified as a compound that activates the mitoUPR, and extends lifespan in an ATFS-1-dependent manner [39]. Combined these results indicate that there are multiple genes or interventions which activate the mitoUPR and extend longevity, but there are also examples in which these phenotypes are uncoupled.

Multiple experiments including the present study have also looked at the effect of the mitoUPR on lifespan directly by either increasing or decreasing the expression of components of the mitoUPR. Knocking down *atfs-1* expression using RNAi does not decrease wild-type lifespan [7, 13, 40], nor do deletions in the *atfs-1* gene decrease wild-type lifespan (**Fig. 6**; [7, 13]). Thus, despite activation of the mitoUPR being correlated with longevity, ATFS-1 is not required for normal lifespan in a wild-type animal.

### ATFS-1 mediates lifespan extension in long-lived mutants

While ATFS-1 is dispensable for wild-type lifespan, ATFS-1 is required for lifespan extension of multiple long-lived mutants. Longevity can be extended by disrupting mito-nuclear protein balance through knocking down the expression of mitochondrial ribosomal protein S5 (*mrsp-5*), which also increases the expression of mitoUPR target gene *hsp-6*. The magnitude of the lifespan extension caused by *mrsp-5* RNAi is decreased by knocking down key mitoUPR component genes *haf-1* or *ubl-5* [11]. In the long-lived mitochondrial mutant *nuo-6*, deletion of *atfs-1* completely reverts the long lifespan to wild-type length, and treatment with *atfs-1* RNAi has similar effects [7]. In mitochondrial mutant *isp-1* worms, knocking down the a key initiatior of mitoUPR, *ubl-5*, decreases their long lifespan but has no effect on the lifespan of wild-type worms [8]. In contrast it has been reported that knockdown of *atfs-1* using RNAi does not decrease *isp-1* lifespan [13]. However, it is possible that in the latter study that the magnitude knockdown was not sufficient to have effects on lifespan, as we and others have found that life-long exposure to *atfs-1* RNAi prevents larval development of *isp-1* worms [7, 41]. Similarly, differing results have been obtained for the requirement of the mitoUPR in the extended lifespan resulting from *cco-1* knockdown. While it has been reported that mutation of *atfs-1* does not decrease lifespan of worms treated with *cco-1* RNAi, despite preventing activation of mitoUPR reporter [13], a subsequent study showed that *atfs-1* RNAi decreases the extent of lifespan extension resulting from *cco-1* RNAi [40]. While differing results have been observed in some cases, overall, these studies suggest a role of ATFS-1 and the mitoUPR in mediating the lifespan extension in a subset of long-lived mutants.

Despite the fact that long-lived mutants with chronic activation of the mitoUPR depend on ATFS-1 for their long lifespan, our current results using the constitutively active *atfs-1(et15)* and *atfs-1(et17)* mutants, as well as previous results using constitutively active *atfs-1* mutants (*et17* and *et18*) show that constitutive activation of ATFS-1 in wild-type worms results in decreased lifespan [13]. This may be partially due to activation of ATFS-1 increasing the proportion of damaged mtDNA when heteroplasmy exists [42]. Consistent with this finding, overexpression of the mitoUPR target gene *hsp-60* also leads to a small decrease in lifespan [43]. In contrast, overexpression of a different mitoUPR target gene, *hsp-6*, is sufficient to increase lifespan [44]. It has also been shown that a hypomorphic reduction-of-function mutation allele of *hsp-6 (mg583)* can also increase lifespan, while *hsp-6* null mutations are thought to be lethal [45]. Combined these results indicate that chronic activation of the mitoUPR is mildly detrimental for lifespan, but that modulation of specific target genes can be beneficial.

### ATFS-1 is necessary for stress resistance in wild-type animals

While ATFS-1 is not required for longevity in wild-type animals, it plays a significant role in protecting animals against exogenous stressors. Disrupting *atfs-1* function decreases organismal resistance to oxidative stress, heat stress, osmotic stress, and anoxia (**Fig. 4**). Additionally, we previously determined that inhibiting *atfs-1* in long-lived *nuo-6* worms completely prevented the increased resistance to oxidative stress, osmotic stress and heat stress typically observed in that mutant [7], and that disruption of *atfs-1* in Parkinson’s disease mutants *pdr-1* and *pink-1* decreased their resistance to oxidative stress, osmotic stress, heat stress, and anoxia [46]. Combined, these results demonstrate that ATFS-1 is required for resistance to multiple types of exogenous stressors.

Even though ATFS-1 is required for the upregulation of stress response genes in response to bacterial pathogens (**Fig. 3**), deletion of *atfs-1* (*gk3094* mutation) did not impact bacterial pathogen resistance. Similarly, another *atfs-1* deletion mutation (*tm4919*) was found not to affect survival during exposure to *P. aeruginosa* [14]. Knocking down *atfs-1* through RNAi inconsistently decreased survival on *P. aeruginosa* (e.g. Fig.3a versus Fig.3h in [14]). While the effect of *atfs-1* disruption on bacterial pathogen resistance was variable, decreasing the expression of a downstream ATFS-1 target gene, *hsp-60*, by RNAi caused a robust decrease in organismal survival on *P. aeruginosa* [43]. As we have previously found that disrupting *atfs-1* induces upregulation of other protective cellular pathways [7] and others have observed a similar phenomenon when a mitoUPR downstream target, *hsp-6*, is disrupted [47], it is possible that the upregulation of other stress pathways may compensate for the inhibition of the mitoUPR, ultimately yielding wild-type or increased levels of resistance to bacterial pathogens, and hiding the normal role of the mitoUPR in resistance to bacterial pathogens.

### Activation of ATFS-1 enhances resistance to exogenous stressors

In this work, we show that constitutive activation of ATFS-1 (*atfs-1(et15)* and *atfs-1(et17)* mutants) is sufficient to increase resistance to multiple different exogenous stressors, including acute oxidative stress, osmotic stress, anoxia and bacterial pathogens. Previous studies have shown that activating the mitoUPR, either through *spg-7* RNAi or through a constitutively active *atfs-1(et15)* mutant, decreased risk of death after anoxia-reperfusion [15], and that constitutively active *atfs-1(et18)* mutants also have increased resistance to *P. aeruginosa* [14]. Overexpression of the mitoUPR target gene *hsp-60* also increases resistance to *P. aeruginosa* [43]. These results support a clear role for ATFS-1 in surviving external stressors.

### ATFS-1 upregulates target genes of multiple stress response pathways

In exploring the mechanism by which ATFS-1 and the mitoUPR modulate stress resistance, we found that activation of ATFS-1, through either a mutation that mildly impairs mitochondrial function (*nuo-6*) or through a mutation that constitutively activates ATFS-1 (*atfs-1(et15)*), causes upregulation of genes involved in multiple stress response pathways including the ER-UPR pathway, the Cyto-UPR pathway, the DAF-16-mediated stress response pathway, the SKN-1-mediated oxidative stress response pathway, the HIF-mediated hypoxia response pathway, the p38-mediated innate immune response pathway and antioxidant genes (**Fig. 2**). These findings are supported by earlier work demonstrating a role for ATFS-1 in upregulating genes involved in innate immunity. Pellegrino *et al*. reported a 16% (59/365 genes) overlap between genes upregulated by activation of the mitoUPR through treatment with *spg-7* RNAi and genes upregulated by exposure a bacterial pathogen [14]. A connection between the mitoUPR and the innate immunity pathway was also suggested by the finding that overexpression of a mitoUPR downstream target, *hsp-60*, increases expression of three innate immunity genes: T24B8.5, C17H12.8 and K08D8.5 [43]. Our results clearly indicate that the role of ATFS-1 in stress response pathways is not limited to the innate immunity, but extends to multiple stress response pathways, thereby providing a mechanistic basis for the effect of ATFS-1 on resistance to stress.

### Conclusions

The mitoUPR is required for animals to survive exposure to exogenous stressors, and activation of this pathway is sufficient to enhance resistance to stress (**Table S4**). In addition to upregulating genes involved in restoring mitochondrial homeostasis, the mitoUPR increases stress resistance by upregulating the target genes of multiple stress response pathways. Although increased stress resistance has been associated with long lifespan, and multiple long-lived mutants exhibit activation of the mitoUPR, constitutive activation of ATFS-1 shortens lifespan while increasing resistance to stress, indicating that the role of ATFS-1 in stress resistance can be experimentally dissociated from its role in longevity. Overall, this work highlights the importance of the mitoUPR in not only protecting organisms from internal stress, but also improving organismal survival upon exposure to external stressors.

## Materials and Methods

### Strains

*C. elegans* strains were obtained from the *Caenorhabditis* Genetics Center (CGC): N2 (wild-type), *nuo-6(qm200)*, *atfs-1(gk3094)*, *nuo-6(qm200);atfs-1(gk3094)*, *atfs-1(et15)*, *atfs-1(et17)*, *ife-2 (ok306)*, *clk-1(qm30)*, *sod-2(ok1030)*, *eat-2(ad1116)*, *osm-5(p813)*, *isp-1(qm150)*, *daf-2(e1370)*, and *glp-1(e2141)*. Strains were maintained at 20°C on nematode grown medium (NGM) plates seeded with OP50 *E. coli*.

### Gene expression in response to stress

#### Stress treatment

Young adult worms were subject to different stress before mRNA was collected. For heat stress, worms were incubated at 35°C for 2 hours and 20°C for 4 hours. For oxidative stress, worms were transferred to plates containing 4 mM paraquat and 100 μM FUdR for 48 hours. For ER stress, worms were transferred to plates containing 5 μg/mL tunicamycin for 24 hours. For osmotic stress, worms were transferred to plates containing 300 mM NaCl for 24 hours. For bacterial pathogen stress, worms were transferred to plates seeded with *Pseudomonas aeruginosa* strain PA14 for 4 hours. For anoxic stress, worms were put in BD Bio-Bag Type A Environmental Chambers (Becton, Dickinson and Company, NJ) for 24 hours and left to recover for 4 hours. For unstressed control conditions, worms were collected at the young adult stage and at Day 1 adult stage. For Day 2 adult control, worms were transferred NGM plates containing 100μM FUdR and collected 2 days later.

#### RNA isolation

RNA was harvested as described previously [48]. Plates of worms were washed three times using M9 buffer to remove bacteria and resuspended in TRIZOL reagent. Worms were frozen in a dry ice/methanol bath and then thawed three times and left at room temperature for 15 minutes. Chloroform was mixed into the tubes and mixture was left to sit at room temperature for 3 minutes. Tubes were then centrifuged at 12,000 g for 15 minutes at 4°C. The upper phase containing the RNA was transferred to a new tube, mixed with isopropanol, and allowed to sit at room temperature for 10 minutes. Tubes were centrifuged at 12,000 g for 10 minutes at 4°C. The RNA pellet was washed with 75% ethanol and resuspended in RNAse-free water.

#### Quantitative RT-PCR

mRNA was converted to cDNA using a High-Capacity cDNA Reverse Transcription kit (Life Technologies/Invitrogen) as described previously [49]. qPCR was performed using a PowerUp SYBR Green Master Mix kit (Applied Biosystems) in a Viia 7 RT-PCR machine from Applied Biosystems. All experiments were performed with least three biological replicates collected from different days. mRNA levels were normalized to act-3 levels and then expressed as a percentage of wild-type. Primer sequences are as follows: *gst-4* (CTGAAGCCAACGACTCCATT, GCGTAAGCTTCTTCCTCTGC), *hsp-4* (CTCGTGGAATCAACCCTGAC, GACTATCGGCAGCGGTAGAG), *hsp-6* (CGCTGGAGATAAGATCATCG, TTCACGAAGTCTCTGCATGG), *hsp-16.2* (CCATCTGAGTCTTCTGAGATTGTT, CTTTCTTTGGCGCTTCAATC), *sod-3* (TACTGCTCGCACTGCTTCAA, CATAGTCTGGGCGGACATTT), *sod-5* (TTCCACAGGACGTTGTTTCC, ACCATGGAACGTCCGATAAC), *nhr-57* (GACTCTGTGTGGAGTGATGGAGAG, GTGGCTCTTGGTGTCAATTTCGGG), *gcs-1* (CCACCAGATGCTCCAGAAAT, TGCATTTTCAAAGTCGGTC), *trx-2* (GTTGATTTCCACGCAGAATG, TGGCGAGAAGAACACTTCCT), *Y9C9A.8* (CGGGGATATAACTGATAGAATGG, CAAACTCTCCAGCTTCCAACA), *T24B8.5* (TACACTGCTTCAGAGTCGTG, CGACAACCACTTCTAACATCTG), *clec-67* (TTTGGCAGTCTACGCTCGTT, CTCCTGGTGTGTCCCATTTT), *dod-22* (TCCAGGATACAGAATACGTACAAGA, GCCGTTGATAGTTTCGGTGT), *ckb-2* (GCATTTATCCGAGACAGCGA, GCTTGCACGTCCAAATCAAC), *act-3* (TGCGACATTGATATCCGTAAGG, GGTGGTTCCTCCGGAAAGAA).

### RNA sequencing and bioinformatic analysis

RNA sequencing was performed previously [50, 51] and raw data is available on NCBI GEO: GSE93724 [51], GSE110984 [7]. Bioinformatic analysis for this study was used to determine differentially expressed genes and identify the degree and significance of overlaps between genes sets.

#### Determining differentially expressed genes

Samples were processed using an RNA-seq pipeline based on the bcbio-nextgen project (https://bcbio-nextgen.readthedocs.org/en/latest/). We examined raw reads for quality issues using FastQC (http://www.bioinformatics.babraham.ac.uk/projects/fastqc/) in order to ensure library generation and sequencing data were suitable for further analysis. If necessary, we used cutadapt http://code.google.com/p/cutadapt/ to trim adapter sequences, contaminant sequences such as polyA tails, and low quality sequences from reads. We aligned trimmed reads to the Ensembl build WBcel235 (release 90) of the *C. elegans* genome using STAR [52]. We assessed quality of alignments by checking for evenness of coverage, rRNA content, genomic context of alignments (for example, alignments in known transcripts and introns), complexity and other quality checks. To quantify expression, we used Salmon [53] to find transcript-level abundance estimates and then collapsed down to the gene-level using the R Bioconductor package tximport [54]. Principal components analysis (PCA) and hierarchical clustering methods were used to validate clustering of samples from the same batches and across different mutants. We used the R Bioconductor package DESeq2 [55] to find differential expression at the gene level. For each wildtype-mutant comparison, we identified significant genes using an FDR threshold of 0.01. Lastly, we included batch as a covariate in the linear model for datasets in which experiments were run across two batches.

#### Venn diagrams

Weighted Venn diagrams were produced by inputting gene lists into BioVenn (https://www.biovenn.nl/). Percentage overlap was determined by dividing the number of genes in common between the two gene sets by the gene list with the smaller gene list.

#### Significance of overlap and enrichment

The significance of overlap between two gene sets was determined by comparing the actual number of overlapping genes to the expected number of overlapping genes based on the sizes of the two gene sets (expected number = number of genes in set 1 X number of genes in set 2/number of genes in genome detected). Enrichment was calculated as the observed number of overlapping genes/the expected number of overlapping genes if genes were chosen randomly.

### Resistance to stress

For acute oxidative stress, young adult worms were transferred onto plates with 300 μM juglone and survival was measured every 2 hours for a total of 10 hours. For chronic oxidative stress, young adult worms were transferred onto plates with 4 mM paraquat and 100 μM FUdR and survival was measured daily until death.

For heat stress, young adult worms were incubated in 37°C and survival was measured every 2 hours for a total of 10 hours. For osmotic stress, young adult worms were transferred to plates containing 450 mM or 500 mM NaCl and survival was measured after 48 hours.

For anoxic stress, plates with young adult worms were put into BD Bio-Bag Type A Environmental Chambers for 75 hours and survival was measured after a 24-hour recovery period.

Two different bacterial pathogenesis assays involving *P. aeruginosa* strain PA14 were performed. In the slow kill assay worms are thought to die from intestinal colonization of the pathogenic bacteria, while in the fast kill assay worms are thought to die from a toxin secreted from the bacteria [25]. The slow kill assay was performed as described previously [14, 27]. In the first protocol [14], PA14 cultures were grown overnight and seeded to center of a 35-mm NGM agar plate. Plates were left to dry overnight, and then incubated in 37°C for 24 hours. Plates were left to adjust to room temperature before approximately 40 L4 worms were transferred onto the plates. The assay was conducted 25°C and plates were checked twice a day until death. In the second protocol [27], overnight PA14 culture were seeded to the center of a 35-mm NGM agar plate containing 20 mg/L FUdR. Plates were incubated at 37°C overnight, then at room temperature overnight before approximately 40 day three adults were transferred onto these plates. The assay was conducted 20°C and plates were checked daily until death. The fast kill pathogenesis assay was performed as described previously [25]. PA14 cultures were grown overnight and seeded to Peptone-Glucose-Sorbitol (PGS) agar plates. Seeded plates were left to dry for 20 minutes at room temperature before incubation at 37°C for 24 hours and then at 23°C for another 24 hours. Approximately 30 L4 worms were transferred onto the plates and were scored as dead or alive at 2, 4, 6, 8 and 24 hours. Fast kill plates were kept at 23°C in between scoring timepoints.

### Lifespan

All lifespan assays were performed at 20°C. Lifespan assays included FUdR to limit the development of progeny and the occurrence of internal hatching. Based on our previous studies, a low concentration of FUdR (25mM) was used to minimize potential effects of FUdR on lifespan [56]. Animals were excluded from the experiment if they crawled off the plate or died of internal hatching of progeny or expulsion of internal organs.

### Statistical Analysis

To ensure unbiased results, all experiments were conducted with the experimenter blinded to the genotype of the worms. For all assays, a minimum of three biological replicates of randomly selected worms from independent populations of worms on different days were used. For analysis of lifespan, oxidative stress, and bacterial pathogen stress, a log-rank test was used. For analysis of heat stress, repeated measures ANOVA was used. For analysis of osmotic stress and anoxic stress, a one-way ANOVA with Dunnett’s multiple comparisons tests was used. For quantitative PCR results we used a two-way ANOVA with Bonferroni post-test. For all bar graphs error bars indicate standard error of the mean and bars indicate the mean.

## Acknowledgments

We would like to thank Paige Rudich for carefully reviewing the manuscript prior to submission. Some strains were provided by the CGC, which is funded by NIH Office of Research Infrastructure Programs (P30 OD010440). We would also like to acknowledge the *C. elegans* knockout consortium and the National Bioresource Project of Japan for providing strains used in this research.

## Author Contributions

Conceptualization: JVR. Methodology: SKS, AT, MM, JVR. Investigation: SKS, AT, MM, JVR. Analysis: SKS, AT, MM, JVR. Visualization: SKS, AT, MM, JVR. Writing – original draft: SKS, JVR. Writing – review and editing: SKS, AT, JVR. Supervision: JVR.

## Materials & Correspondence

Correspondence and material requests should be addressed to Jeremy Van Raamsdonk.

## Data availability

RNA-seq data has been deposited on GEO: GSE93724, GSE110984. All other data and strains generated in the current study are included with the manuscript or available from the corresponding author on request.

## Supplementary Figures

**Figure S1.**
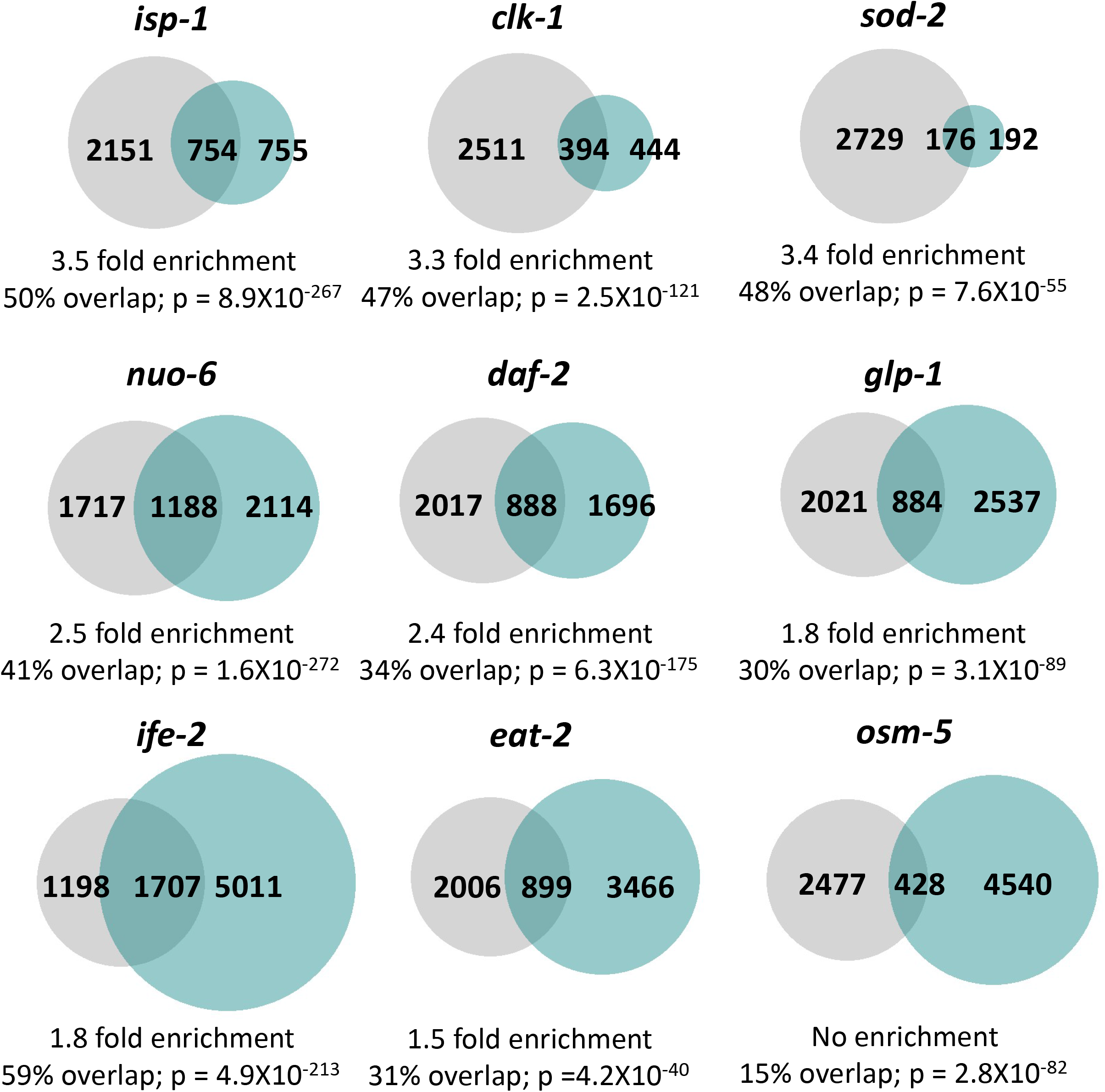
Multiple long-lived mutants from different pathways of lifespan extension show upregulation of ATFS-1-dependent genes. To determine the extent to which long-lived genetic mutants from different pathways of lifespan extension show differential expression of ATFS-1 target genes, we compared genes that are upregulated in nine different long-lived mutants to genes upregulated in a constitutively active *atfs-1* mutant (*et15*). All of the long-lived mutant worms, except for *osm-5*, show a highly significant degree of overlap with the constitutively active *atfs-1* mutant. The grey circles represent genes that are significantly upregulated in the constitutively active *atfs-1(et15)* mutant. Turquoise circles are genes that are significantly upregulated in the long-lived mutant indicated. The number of unique and overlapping genes are indicated. Overlap is calculated as the number of genes in common between the two gene sets divided by the smaller gene set. Enrichment is calculated as the number of overlapping genes observed divided by the number of overlapping genes predicted if genes were chosen randomly.

**Table S1.** Target genes examined for each stress response pathway.

| Target Gene | Stress response pathway |
| --- | --- |
| *hsp-6* | Mitochondrial unfolded protein response |
| *hsp-4* | ER unfolded protein response |
| *hsp-16.2* | Cytoplasmic unfolded protein response |
| *sod-3* | DAF-16-mediated stress response pathway |
| *gst-4* | SKN-1-mediated stress response pathway |
| *nhr-57* | HIF-1-mediated hypoxia pathway |
| *Y9C9A.8* | p38-mediated innate immune pathway |
| *trx-2* | Antioxidant genes |

**Table S4.** Effect of modulating ATFS-1 levels and activation on stress resistance, lifespan and expression of stress response genes.

|  | *atfs-1(et15)*<br>Gain-of-function | *atfs-1(et17)*<br>Gain-of-function | *atfs-1(gk3094)*<br>Loss-of-function |
| --- | --- | --- | --- |
| Acute oxidative stress resistance | ↑ | ↑ | ↓ |
| Chronic oxidative stress resistance | ↓ | ↑ | ↓ |
| Heat stress resistance | ↓ | = | ↓ |
| Osmotic stress resistance | ↑ | ↑ | ↓ |
| Anoxia resistance | ↑ | ↑ | ↓ |
| Bacterial pathogens resistance | == | ↓ ↑ | ↑ ↑ |
| Lifespan | ↓ | ↓ | = |
| *hsp-6* expression (mitoUPR) | ↑ | ↑ | = |
| *hsp-4* expression (ER-UPR) | = | = | = |
| *hsp-16.2* expression (Cyto-UPR) | ↑ (NS) | ↑ (NS) | ↑ (NS) |
| *sod-3* expression (DAF-16) | ↑ | ↑ | = |
| *gst-4* expression (SKN-1) | = | = | = |
| *nhr-57* expression (HIF-1) | ↑ | = | = |
| *Y9C9A.8* expression (Innate immunity) | ↑ | ↑ | = |
| *trx-2* (antioxidant) | ↑ | ↑ | = |

